# Neuroligin-3 in Dopaminergic Circuits Promotes Behavioral and Neurobiological Adaptations to Chronic Morphine Exposure

**DOI:** 10.1101/2022.03.14.484267

**Authors:** Dieter D. Brandner, Cassandra L. Retzlaff, Adrina Kocharian, Bethany J. Stieve, Mohammed A. Mashal, Paul G. Mermelstein, Patrick E. Rothwell

## Abstract

Chronic opioid exposure causes structural and functional changes in brain circuits, which may contribute to opioid use disorders. Synaptic cell-adhesion molecules are prime candidates for mediating this opioid-evoked plasticity. Neuroligin-3 (NL3) is an X-linked postsynaptic adhesion protein that shapes synaptic function at multiple sites in the mesolimbic dopamine system. We therefore studied how genetic knockout of NL3 alters responses to chronic morphine in male mice. Constitutive NL3 knockout caused a persistent reduction in psychomotor sensitization after chronic morphine exposure and changed in the topography of locomotor stimulation produced by morphine. This latter change was recapitulated by conditional genetic deletion of NL3 from cells expressing the Drd1 dopamine receptor, whereas reduced psychomotor sensitization was recapitulated by conditional genetic deletion from dopamine neurons. Without NL3 expression, dopamine neurons in the ventral tegmental area exhibited diminished activation following chronic morphine exposure, by measuring *in vivo* calcium signals with fiber photometry. This altered pattern of dopamine neuron activity may be driven by aberrant forms of opioid-evoked synaptic plasticity in the absence of NL3: dopamine neurons lacking NL3 showed weaker synaptic inhibition at baseline, which was subsequently strengthened after chronic morphine. In total, our study highlights neurobiological adaptations in dopamine neurons of the ventral tegmental area that correspond with increased behavioral sensitivity to opioids, and further suggests that NL3 expression by dopamine neurons provides a molecular substrate for opioid-evoked adaptations in brain function and behavior.

## INTRODUCTION

Drugs of abuse target the mesolimbic dopamine projection, with a final common pharmacologic effect of increased extracellular dopamine levels in the nucleus accumbens (NAc)^1^. Morphine and other opioids specifically increase the firing rate of dopamine neurons in the ventral tegmental area (VTA) by activating mu opioid receptors expressed on inhibitory synaptic inputs^2^, thereby enhancing dopamine release via disinhibition. Chronic opioid exposure produces various neurobiological adaptations in mesolimbic circuitry, including functional and structural changes at synaptic connections in both the VTA and NAc^3,4^. These adaptations may contribute to the development of opioid use disorders, but their molecular basis remains poorly understood.

Synaptic cell adhesion molecules play a critical role in sculpting synaptic structure and function^5^, and thus represent potential molecular mediators of drug-evoked plasticity^6^. This includes the neurexin and neuroligin families of adhesion molecules, which have been genetically linked to substance use disorders and other neuropsychiatric conditions^7-10^. Neuroligin-3 (NL3) is a particularly intriguing candidate for mediating drug-evoked plasticity within the mesolimbic dopamine system, since it has functional roles in both VTA dopamine neurons and NAc medium spiny projection neurons (MSNs). Genetic knockout of NL3 alters the subunit composition of AMPA receptors in VTA dopamine neurons^11^ and causes synaptic disinhibition of D1-MSNs in the NAc^12^. Similar forms of synaptic dysfunction have been reported after chronic exposure to drugs of abuse^13-15^, suggesting genetic knockout of NL3 may alter behavioral and neurobiological responses to addictive substances like morphine.

To explore the potential role of NL3 in neurobehavioral adaptations caused by chronic morphine exposure, we studied the impact of constitutive genetic knockout of NL3 and conditional genetic knockout from specific cell types within mesolimbic circuits. We report that constitutive NL3 knockout males exhibit a change in spatial organization of locomotor activity following morphine exposure, which we refer to as locomotor topography^16^, and a persistent attenuation of psychomotor sensitization following chronic morphine exposure^17^. Conditional genetic knockout revealed a double-dissociation in the cell types where NL3 deletion produced these effects: NL3 deletion from D1-MSNs altered locomotor topography after morphine exposure, whereas NL3 deletion from dopamine neurons persistently attenuated psychomotor sensitization following chronic morphine.

Psychomotor sensitization is one behavioral manifestation of the enhanced sensitivity of the mesolimbic dopamine system that develops after intermittent opioid exposure^18^, and is initiated by opioid receptor activation in the VTA^19,20.^ We therefore explored morphine-evoked neurobiological adaptations in dopamine neurons. In the absence of NL3, VTA dopamine neurons showed an attenuation of activity after chronic morphine exposure and altered inhibitory synaptic input. Together, these results indicate that NL3 has multiple distinct functions integral to the behavioral opioid response at discrete loci in the mesolimbic dopamine system. The function of NL3 in VTA dopamine neurons appears to be particularly critical for regulating their adaptations to chronic morphine exposure, and for long-lasting enhancement of behavioral sensitivity to opioids.

## MATERIALS AND METHODS

### Subjects

Mice were housed in groups of 2-5 per cage, on a 12-hour light cycle (0600h–1800h) at ∼23° C with ad libitum food and water access. Experimental procedures were conducted between 1000h–1600h and were approved by to the University of Minnesota Institutional Animal Care and Use Committee and observed the NIH *Guidelines for the Care and Use of Laboratory Animals*. All mouse lines were maintained on a C57Bl/6J genetic background. Constitutive NL3 knockout animals (The Jackson Laboratory Strain #008394) were generated as previously described^21^. Conditional NL3 knockout animals were generated as previously described^12^, with loxP sites flanking exons 2 and 3 of the *Nlgn3* gene. Floxed NL3 mice were crossed with Drd1-Cre BAC transgenic FK150^22^, Adora2a-Cre BAC transgenic KG139^23^, or DAT-IRES-Cre knock-in^24^. These Cre driver lines were used to conditionally knockout NL3 in D1-MSNs, D2-MSNs, or dopamine neurons, respectively. Due to X-linkage of the *Nlgn3* gene, experiments were conducted using male mice since it was possible to generate wild-type and knockout littermates from a single breeding scheme and thus control for parental genotype.

### Morphine Administration

Morphine hydrochloride (Mallinckrodt) was dissolved in sterile 0.9% saline and delivered subcutaneously (5 mL/kg) by bolus injection at one of several doses (2, 6.32, or 20 mg/kg). We have previously used these doses to measure the acute psychomotor response to morphine^25^ and long-lasting psychomotor sensitization^26^. The day prior to the first acute morphine injection, animals were habituated to the behavior apparatus and given an equivalent volume of 0.9% saline subcutaneously. This was followed by exposure to increasing morphine doses of 2, 6.32, and 20 mg/kg morphine on consecutive days. Psychomotor sensitization consisted of seven consecutive doses of daily morphine administered subcutaneously at 20 mg/kg. To minimize the contextual effects of the behavior apparatus on psychomotor sensitization, locomotion was only measured during saline habituation and after morphine administration on Day 1 and Day 7. Morphine challenge injections were delivered 21 days following the final day of morphine sensitization and were conducted in a manner analogous to acute dose-response.

### Fiber Photometry

Continuous fiber photometry recordings were conducted for 15 minutes prior to and 30 minutes following morphine injection on Days 1 and 7. During recordings, mice were placed in an empty home cage, to closely monitor torsion on the patch cord generated by locomotion after morphine exposure. Real-time data were acquired using a RZ5P fiber photometry workstation (Tucker Davis Technologies). As previously described^27^, 470 nm and 405 nm LEDs (ThorLabs) were modulated at distinct carrier frequencies (531 Hz and 211 Hz, respectively), and passed through a fluorescence mini cube (Doric Lenses) coupled into a patch cord (400 μm, 0.48 NA). The distal end of the patch cord was connected to the implanted fiberoptic cannula by a fitted ceramic sleeve. Fluorescence was projected back through the mini cube and focused onto a photoreceiver (Newport Model 2151). Signals were sampled at 6.1 kHz, demodulated in real-time, and saved for offline analysis. The change in fluorescent signal (ΔF/F) was calculated within each session as described^28^. Each channel was low-passed filtered (<2Hz) and a linear least-squared model fit the isosbestic control signal (405 nm) to the calcium-dependent signal (470 nm). Data was smoothed by calculating the mean ΔF/F over a sliding window of 50 sec along the entire duration of the normalized ΔF/F recording, to reveal slow fluctuations in GCaMP fluorescence caused by morphine exposure. Average fluorescent signal for 200 sec prior to morphine injection and 1000 sec following morphine injection was computed to determine the pre-morphine baseline and post-morphine ΔF/F. The difference between these two averages was calculated as an index of VTA dopamine neuron activity in response to morphine administration. After the end of repeated morphine administration, the D2 agonist apomorphine (5 mg/kg, s.c.) was administered on Day 8. Recordings were analyzed as indicated above to compute average ΔF/F fluorescence before and after injection. Groups exhibited an average drop in ΔF/F immediately following administration of apomorphine due to activation of somatodendritic D2 autoreceptors on VTA dopamine neurons^29^.

### Electrophysiology

NL3^fl/y^;DAT^Cre/wt^ and NL3^wt/y^;DAT^Cre/wt^ animals that received stereotaxic injection of AAV9-EF1a-DIO-eYFP were allowed to recover 5-14 days before the start of drug treatment. All mice were habituated to injection and then treated with either morphine (20 mg/kg, sc) or saline for seven consecutive days. Twenty-four hours after the final injection, mice were deeply anesthetized with isoflurane and perfused with ice cold sucrose solution containing (in mM): 228 sucrose, 26 NaHCO3, 11 glucose, 2.5 KCl, 1 NaH2PO4-H2O, 7 MgSO4-7H20, 0.5 CaCl2-2H2O. Brains were then rapidly dissected and placed in cold sucrose for slicing. Horizontal hemisected slices (225 μm) containing the VTA were collected using a vibratome (Leica VT1000S) and allowed to recover in a warm (33°C) holding chamber with artificial cerebrospinal fluid (aCSF) containing (in mM): 119 NaCl, 26.2 NaHCO3, 2.5 KCl, 1 NaH2PO4-H2O, 11 glucose, 1.3 MgSO4-7H2O, 2.5 CaCl2-2H2O. Slices recovered for 10-15 min and then equilibrated to room temperature for at least one hour before use. Slices were transferred to a submerged recording chamber and continuously perfused with aCSF at a rate of 2 mL/min at room temperature. All solutions were continuously oxygenated (95% O2/5% CO2).

Dopamine neurons in the VTA were identified by viral expression of eYFP using an Olympus BX51W1 microscope. Whole-cell voltage-clamp recordings were made with borosilicate glass electrodes (3-5 MΩ) filled with (in mM): 125 CsCl, 10 TEA-Cl, 10 HEPES, 0.1 EGTA, 3.3 QX-314 (Cl^-^ salt), 1.8 MgCl2, 4 Na2-ATP, 0.3 Na-GTP, 8 Na2-phosphocreatine (pH 7.3 adjusted with CsOH; 276 mOsm). All recordings were performed using a MultiClamp 700B amplifier (Molecular Devices), filtered at 2 kHz, and digitized at 10 kHz. Data acquisition and analysis were performed online using Axograph software. Series resistance was monitored continuously, and experiments were discarded if resistance changed by >20%. To pharmacologically isolate spontaneous inhibitory postsynaptic currents, the extracellular bath solution contained the AMPA receptor antagonist NBQX (10 uM) and the NMDA receptor antagonist D-APV (50µM). These were inward currents at a holding potential of −60 mV, due to the high chloride concentration in the internal pipette solution. At least 200 events per cell analyzed was acquired across 15 sec sweeps, filtered at 0.5 kHz, and detected using an amplitude threshold of 5.5-6.5 pA and a signal-to-noise ratio threshold of 2.5 standard deviations.

### Experimental Design and Statistical Analyses

Constitutive NL3 knockout mice (Figure 1) were generated by breeding heterozygous females (NL3^-/+^) with wild-type males (NL3^+/y^). Conditional NL3 knockout mice were generated by crossing NL3^fl/fl^ dams with sires that were Drd1-Cre hemizygotes (Figure 2), Adora2a-Cre hemizygotes (Figure 3), or DAT^Cre/wt^ heterozygotes (Figure 4). For fiber photometry experiments (Figure 5) and slice physiology experiments (Figure 6), homozygous DAT^Cre/Cre^ sires were crossed with heterozygous NL3^fl/+^ dams, so that all offspring were DAT^Cre/wt^ heterozygotes with or without a floxed NL3 allele. These breeding schemes generated male offspring with either intact or complete loss of NL3 expression, but female offspring with only partial loss of NL3 expression, so we focused our analysis on male mice in this manuscript. For clarity, the details regarding each of these breeding schemes are repeated at the beginning of each section describing the corresponding results.

**Figure 1.**
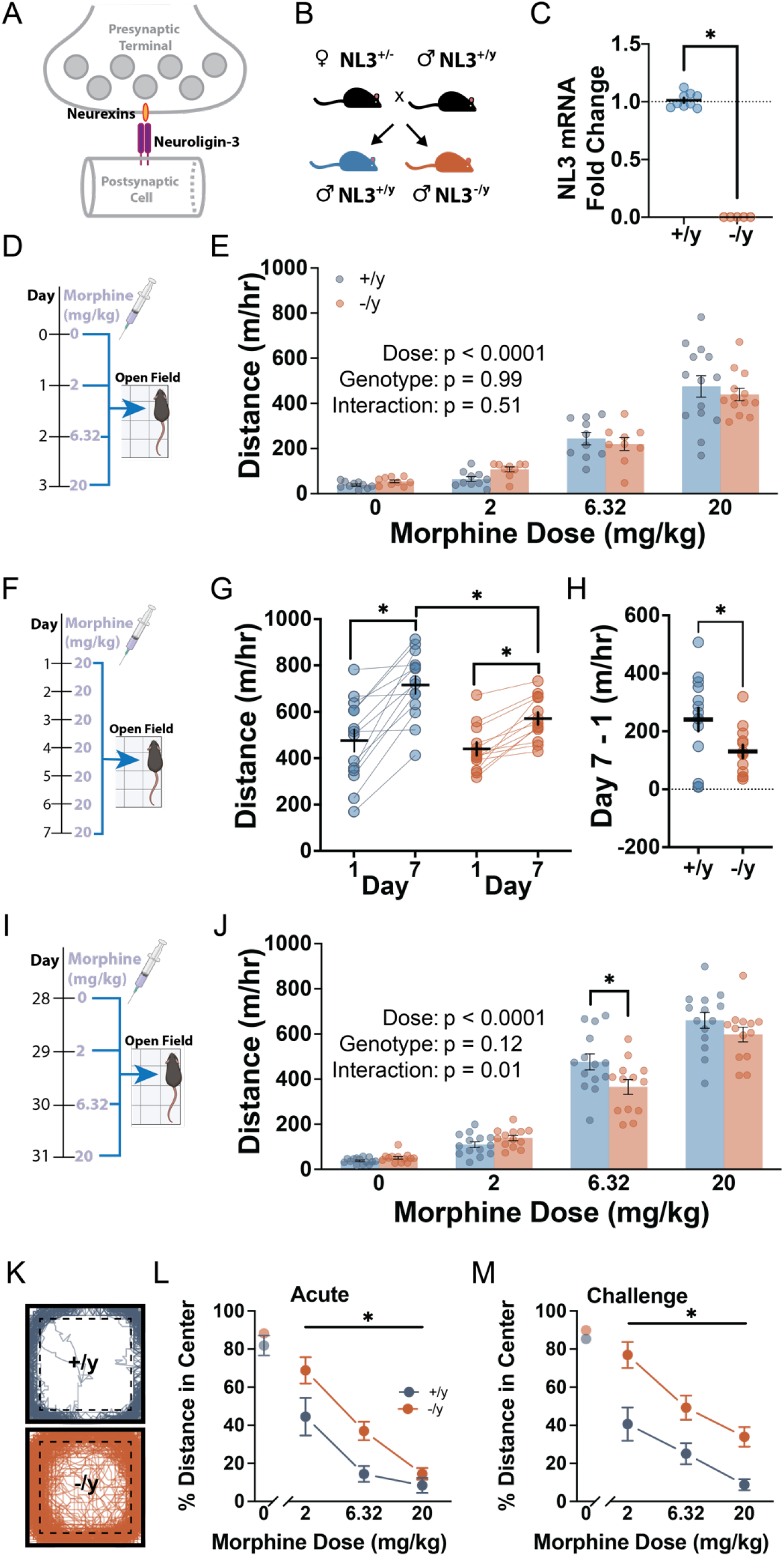
Responses to acute and chronic morphine exposure in NL3 constitutive knockout mice. **(A)** Schematic showing synaptic localization of NL3. **(B)** Breeding scheme. **(C)** Quantification of NL3 mRNA in striatal tissue using quantitative RT-PCR in +/y (n=9) and - /y (n=5) littermates. **(D)** Schematic showing dose sequence for acute morphine exposure. **(E)** Distance travelled following each acute injection in +/y (n=10) and -/y (n=9) littermates at 0-6.32 mg/kg, and +/y (n= 14) and - /y (n=13) littermates at 20 mg/kg, along with statistical results from factorial ANOVA analysis. **(F)** Schematic showing chronic morphine exposure with locomotor activity tested on Days 1 and 7. **(G)** Distance travelled following morphine exposure on Days 1 and 7 in +/y (n=14) and -/y (n=13) littermates. **(H)** Difference in locomotion on Days 1 and 7 in the same cohort of animals. **(I)** Schematic showing dose sequence for morphine challenge. **(J)** Distance travelled following each challenge injection of morphine, along with statistical results from factorial ANOVA analysis. **(K)** Track plots illustrating path of travel after challenge with 6.32 mg/kg morphine. **(L, M)** Percentage of total distance traveled in the central area after acute (**L**) or chronic (**M**) morphine exposure, in the same groups shown above. *p<0.05 comparing groups with simple effect tests **(G)**, unpaired t-test **(H)**, Fisher’s LSD *post-hoc* test **(J)**, or main effect in ANOVA **(L, M)**.

**Figure 2.**
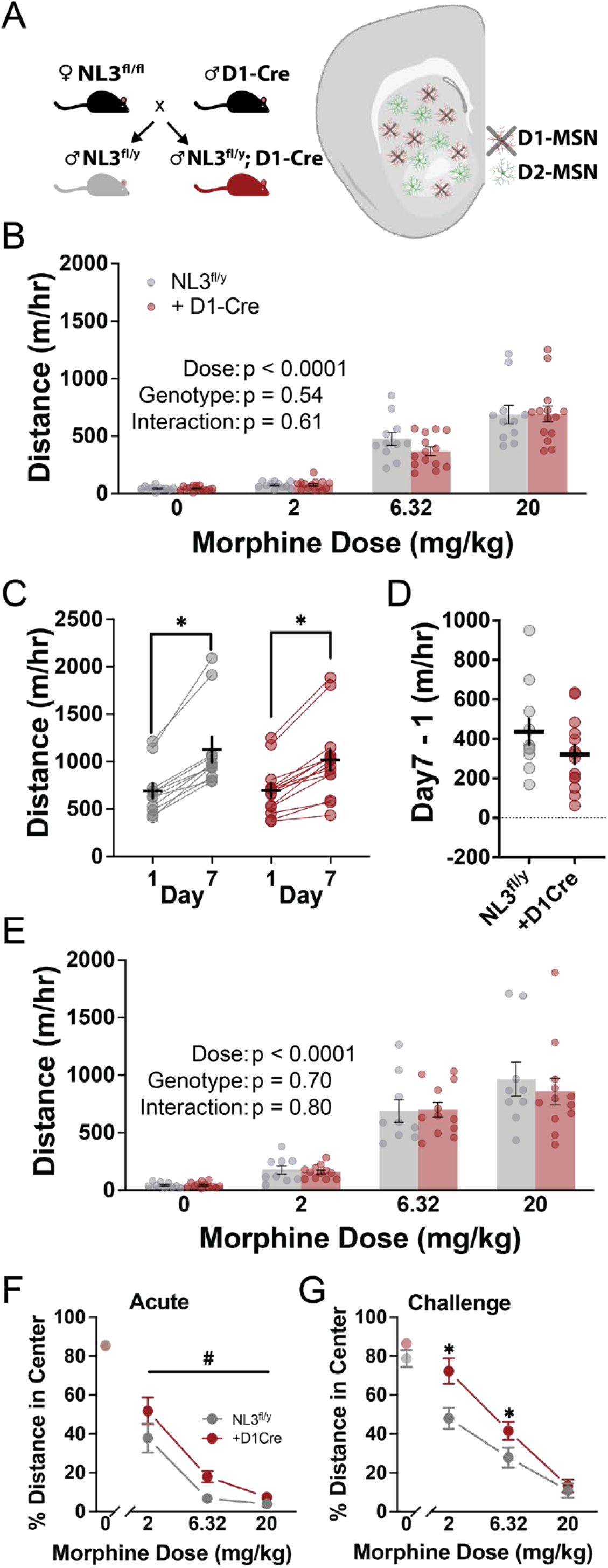
Responses to acute and chronic morphine exposure after conditional NL3 knockout from D1-MSNs. **(A)** Breeding scheme used to achieve conditional knockout of NL3 in D1-MSNs. **(B)** Distance travelled following each acute injection in NL3^fl/y^, without (n=11) and with D1-Cre (n=14), along with statistical results from factorial ANOVA. **(C)** Distance travelled for the same cohorts following chronic morphine exposure on Days 1 and 7. **(D)** Difference in locomotion on Days 1 and 7. **(E)** Distance travelled following each challenge injection of morphine, along with statistical results from factorial ANOVA analysis. **(F, G)** Percentage of total distance traveled in the central area after acute (**F**) or chronic (**G**) morphine exposure, in the same groups shown above. *p<0.05 and #p<0.10 for simple effect of Day in each genotype **(C)**, main effect in ANOVA **(F)**, or Fisher’s LSD *post-hoc* test **(G)**.

**Figure 3.**
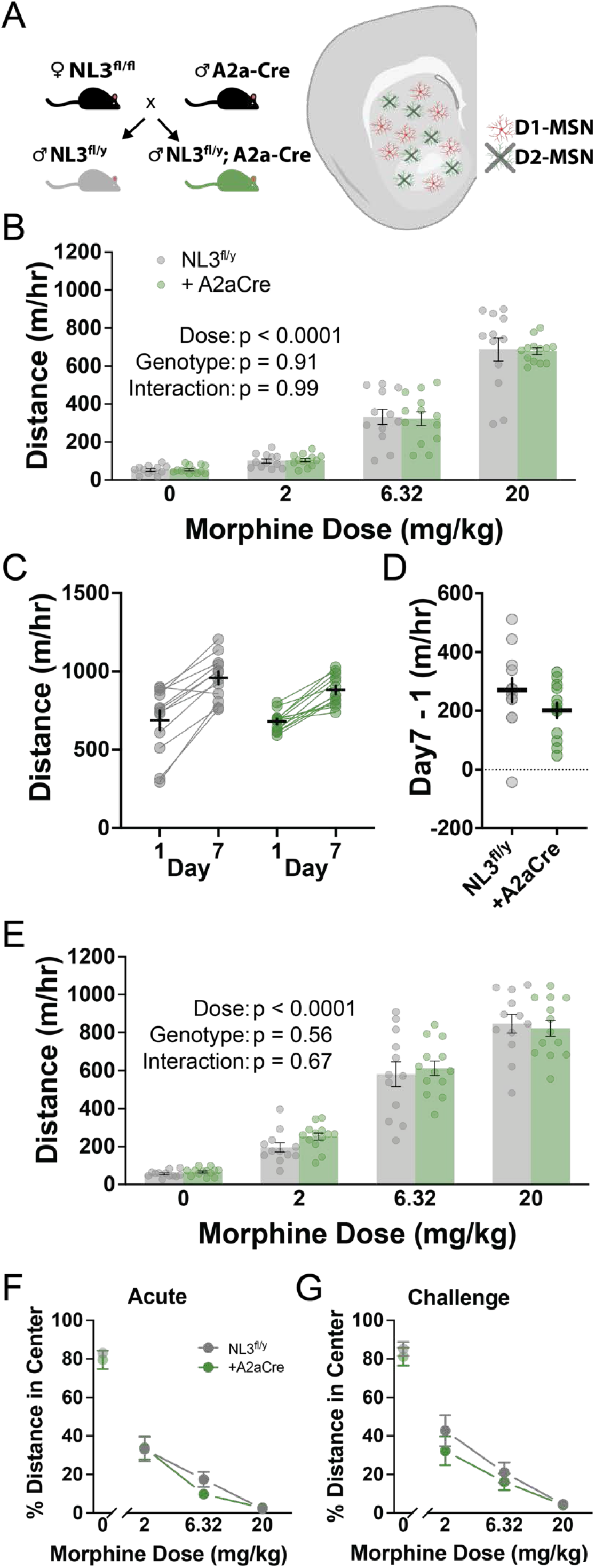
Responses to acute and chronic morphine exposure after conditional NL3 knockout from D2-MSNs. **(A)** Breeding scheme used to achieve conditional knockout of NL3 in D1-MSNs. **(B)** Distance travelled following each acute injection in NL3^fl/y^, without (n=12) and with A2a-Cre (n=13), along with statistical results from factorial ANOVA. **(C)** Distance travelled for the same cohorts following chronic morphine exposure on Days 1 and 7. **(D)** Difference in locomotion on Days 1 and 7. **(E)** Distance travelled following each challenge injection of morphine, along with statistical results from factorial ANOVA analysis. **(F, G)** Percentage of total distance traveled in the central area after acute (**F**) or chronic morphine exposure, in the same groups shown above. *p<0.05 for simple effect of Day in each genotype **(C)**.

**Figure 4.**
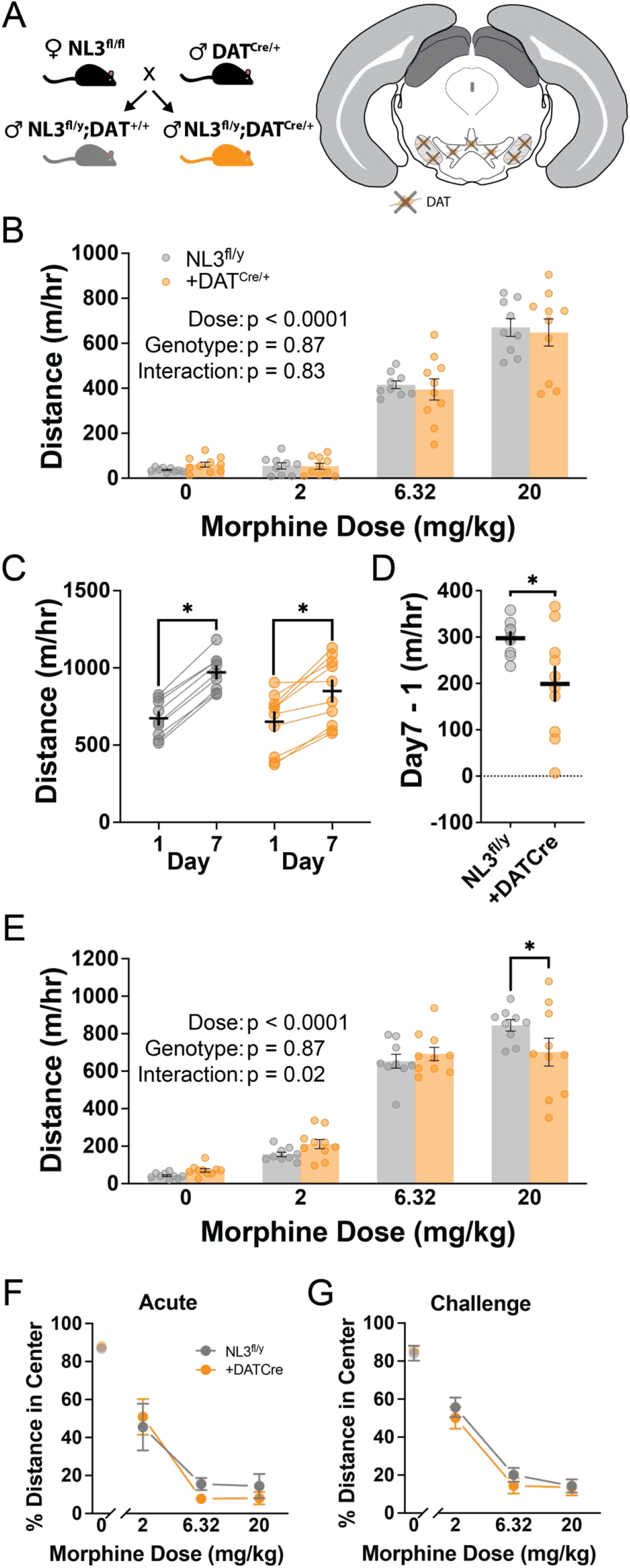
Responses to acute and chronic morphine exposure after conditional NL3 knockout from dopamine neurons. **(A)** Breeding scheme used to achieve conditional knockout of NL3 in dopamine neurons. **(B)** Distance travelled following each acute injection in NL3^fl/y^, without (n=9) and with DAT-Cre (n=10), along with statistical results from factorial ANOVA. **(C)** Distance travelled for the same cohorts following chronic morphine exposure on Days 1 and 7. **(D)** Difference in locomotion on Days 1 and 7. **(E)** Distance travelled following each challenge injection of morphine, along with statistical results from factorial ANOVA analysis. **(F, G)** Percentage of total distance traveled in the central area after acute (**F**) or chronic (**G**) morphine exposure, in the same groups shown above. *p<0.05 for simple effect of Day in each genotype **(C)**, unpaired t-test **(D)**, or Fisher’s LSD *post-hoc* test **(E)**.

**Figure 5.**
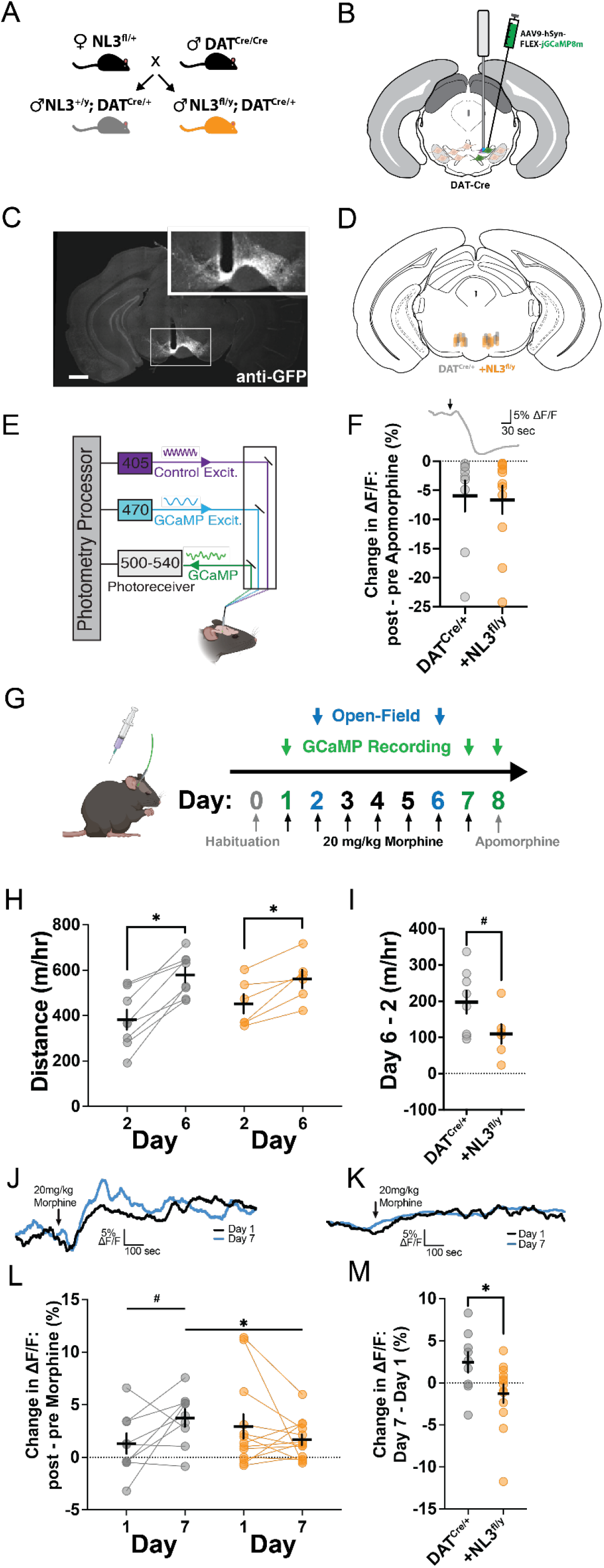
Fiber photometry recordings from VTA dopamine neurons after chronic morphine exposure. **(A)** Breeding scheme used to generate DAT-Cre mice with or without floxed NL3. **(B)** Cartoon showing unilateral stereotaxic injection of AAV9-hSyn-FLEX-jGCaMP8m into the VTA, with fiber optic implant placed just above the injection site. (C) Representative image of jGCaMP8m expression in the VTA (Scale bar = 500µm). **(D)** Anatomical locations of fiber optic implants in individual mice of each genotype. **(E)** Cartoon showing fiber photometry system. **(F)** Effect of apomorphine (5mg/kg, s.c.), injected at the time indicated by the arrow (top), on fluorescent signal in DAT^Cre/+^ mice without (n=9) or with NL3^fl/y^ (n=13). **(G)** Experimental timeline showing habituation (Day 0), 20 mg/kg morphine injections (Days 1-7), and apomorphine injection (Day 8). Photometry recordings were performed on Day 1 and Day 7, and locomotion was tested on Day 2 and Day 6. **(H)** Distance travelled following morphine exposure on Days 2 and 6, measured in a subset of DAT^Cre/+^ mice without (n=8) or with NL3^fl/y^ (n=6). **(I)** Difference in locomotion on Days 1 and 7. **(J, K)** Representative photometry traces from DAT^Cre/+^ mice without **(J)** or with NL3^fl/y^ **(K)**, on Day 1 and Day 7 of morphine exposure. **(L)** Difference in fluorescent signal before and after morphine injection on Day 1 and Day 7. **(M)** Change in difference score across days for individual mice. *p<0.05 or #p<0.10 for simple effect of Day in each genotype **(H)**, unpaired t-test **(I, M)**, or Fisher’s LSD *post hoc* test **(L)**.

**Figure 6.**
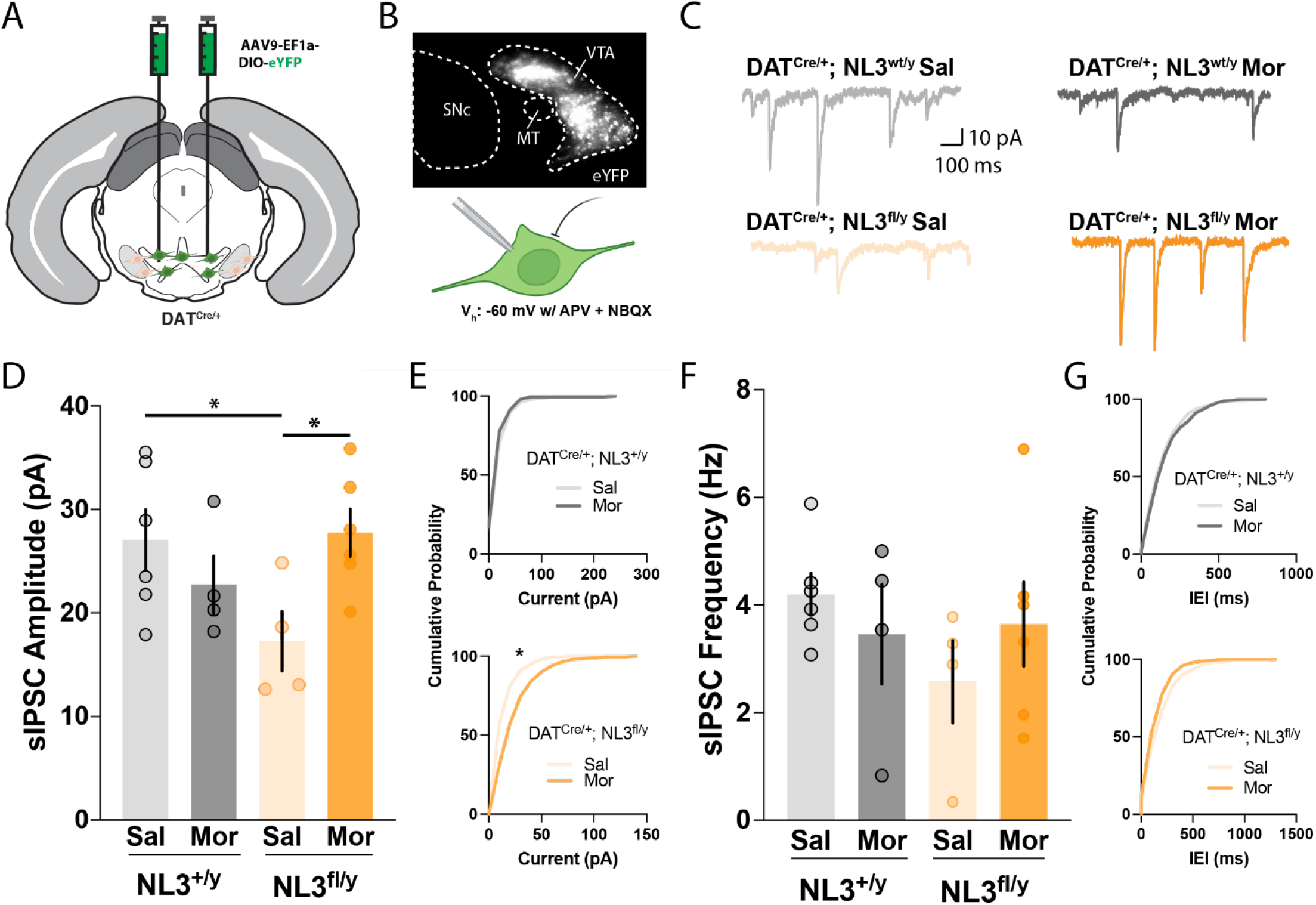
Synaptic inhibition onto VTA dopamine neurons before and after morphine. **(A)** Cartoon showing bilateral stereotaxic injection of AAV9-EF1a-DIO-eYFP virus into the VTA. **(B)** Fluorescent micrograph (top) shows VTA dopamine neurons expressing Cre-dependent YFP in relation to the medial terminal nucleus of the accessory optic tract (MT) and the substantia nigra pars compacta (SNC). Cartoon (bottom) shows recording of spontaneous inhibitory postsynaptic currents (sIPSCs) at a holding potential of −60 mV, in the presence APV and NBQX. **(C)** Example traces from each genotype. **(D)** Average sIPSC amplitude in cells from each group (NL3^+/y^ Sal n=6 cells from 3 animals; NL3^+/y^ Mor n=4 cells from 3 animals; NL3^fl/y^ Sal n=4 cells from 3 animals; NL3^fl/y^ Mor n=6 cells from 3 animals). **(E)** Cumulative probability plots for sIPSC amplitude in NL3^+/y^ (top) and NL3^fl/y^ (bottom). **(F)** Average sIPSC frequency in cells from each group. **(G)** Cumulative probability plots for sIPSC inter-even interval (IEI) in NL3^+/y^ (top) and NL3^fl/y^ (bottom). *p<0.05 comparing groups with Fisher’s LSD *post hoc* test (D) or the Kolmogorov-Smirnov test **(E)**.

Sample sizes are indicated in figure legends and graphs that display individual data points, along with measures of central tendency (mean) and variability (SEM). Data were analyzed in factorial ANOVA models using GraphPad Prism 9 and IBM SPSS Statistics v24, with repeated measures on within-subject factors or mixed-effect models to handle missing data. For main effects or interactions involving repeated measures, the Greenhouse-Geisser correction was applied to control for potential violations of the sphericity assumption. This correction reduces the degrees of freedom, resulting in non-integer values. Significant interactions were decomposed by analyzing simple effects (i.e., the effect of one variable at each level of the other variable). Significant main effects were analyzed using LSD post-hoc tests. The Type I error rate was set to α=0.05 (two-tailed) for all comparisons

## RESULTS

### NL3 knockout mice exhibit less psychomotor sensitization after chronic morphine exposure

To study the effects of genetically deleting NL3 from all synapses in the brain (Figure 1A), we utilized a constitutive NL3 knockout mouse line^21^. Since the gene that encodes NL3 is located on the X-chromosome, we generated male NL3 hemizygous knockouts (NL3^-/y^) and wild-type littermates (NL3^+/y^) by breeding heterozygous females (NL3^-/+^) with wild-type males (NL3^+/y^) (Figure 1B). Genetic knockout was validated using pooled bilateral tissue punches from the dorsal striatum and nucleus accumbens (Figure 1C), which demonstrated robust depletion of NL3 mRNA in knockout mice (t_12_=35.70, p<0.0001). We first examined behavioral response to acute morphine exposure by injecting increasing doses of morphine (2-20 mg/kg, s.c.) prior to a test of open field activity (Figure 1D). All mice exhibited a dose-dependent increase in locomotor activity (Figure 1E), as indicated by a main effect of Dose (F_1.45,36.87_=90.23, p<0.0001). However, there was no main effect of Genotype (F_1,76_<1) or interaction (F_3,76_<1), indicating that distance traveled was similar in both genotypes after acute morphine exposure.

We next measured behavioral responses following seven daily injections of 20 mg/kg morphine (Figure 1F), a chronic pattern of exposure that causes psychomotor sensitization alongside analgesic tolerance^26^. Open field activity was measured after injection on Days 1 and 7 (Figure 1G), to monitor the development of psychomotor sensitization. As expected, distance traveled increased substantially after chronic morphine administration, as indicated by a main effect of Day (F_1,25_=61.79, p<0.0001). There was also a significant Day x Genotype interaction (F_1,25_=5.40, p=0.029), indicating a significant reduction in distance travelled by NL3^-/y^ mutants versus NL3^+/y^ controls on Day 7 (p=0.014), despite similar distance traveled on Day 1. This result was most apparent after computing the change in distance traveled between Day 1 and Day 7 for each individual animal (Figure 1H), revealing a significant reduction in psychomotor sensitization in NL3^-/y^ mutants (t_25_=2.32, p=0.029). Notably, the antinociceptive effect of acute morphine exposure and development of antinociceptive tolerance were similar in NL3^-/y^ and NL3^+/y^ mice (Table S1), indicating that NL3 plays a role in some but not all behavioral adaptations to chronic morphine exposure.

We have previously documented the durability of psychomotor sensitization after this regimen of chronic morphine exposure^26^. We thus waited 21 days after the end of chronic exposure, and then challenged the same cohorts of animals with increasing doses of morphine (2-20 mg/kg) (Figure 1I). Analysis of distance travelled in the open field (Figure 1J) revealed a significant Dose × Genotype interaction (F_3,75_=3.886, p=0.012), a phenotype not observed after acute morphine exposure (Figure 1E). Distance travelled after a challenge injection of 6.32 mg/kg morphine was significantly reduced in NL3^-/y^ mutants versus NL3^+/y^ controls (p=0.030). These results suggest that neuroadaptations supporting psychomotor sensitization are dependent upon NL3 for complete and persistent expression.

### Altered locomotor topography after morphine exposure in NL3 knockout mice

After morphine injection, the locomotor hyperactivity exhibited by wild-type mice leads to a path of travel along the perimeter of the activity chamber^16^. In contrast, NL3^-/y^ mutants exhibited a more curvilinear path of travel after morphine injection, bypassing the corners of the chamber and instead traveling through more central areas (Figure 1K). We quantified the topography of locomotor activity by calculating the percentage of total distance traveled through the interior of the chamber. There was no significant difference between genotypes in this measure prior to acute morphine exposure (Figure 1L) (t_14_=1.13, p=0.28), and a non-significant trend prior to morphine challenge (Figure 1M) (t_22_=1.98, p=0.06). There was a main effect of Dose after both acute morphine exposure (F_1.47,20.63_=46.67, p<0.001) and morphine challenge (F_1.69,35.57_=35.59, p<0.001), indicating that higher doses of morphine caused a greater shift in the distribution of locomotor activity to the perimeter of the activity chamber. There was also a main effect of Genotype after both acute morphine exposure (F_1,14_=7.81, p=0.014) and morphine challenge (F_1,21_=18.78, p<0.001), supporting the conclusion that NL3^-/y^ mutants have a greater percentage of distance travelled through the interior of the chamber. After morphine administration, the sharper turns made by NL3^-/y^ mutants to bypass the corners of the chamber were also reflected by significantly higher absolute angular acceleration (Table S2), while there were no differences between genotypes in the spatial scaling exponent^30^ (Table S3). Changes in locomotor topography were evident after both acute and chronic morphine exposure, whereas distance travelled by NL3^-/y^ mutants was only different after chronic morphine exposure, providing initial evidence that these two phenotypes may be dissociable.

### Conditional NL3 deletion from D1-MSNs alters locomotor topography but not sensitization after morphine exposure

We next sought to determine if behavioral responses to morphine are regulated by conditional NL3 deletion from specific cell types within the mesolimbic dopamine system. Prior work has shown that NL3 is expressed at a particularly high level by D1-MSNs in the NAc, and conditional genetic deletion from this specific cell type recapitulates some behavioral phenotypes of constitutive NL3 knockout mice^12^. Synaptic inhibition of D1-MSNs is diminished in the absence of NL3 expression^12^, and this D1-MSN disinhibition may drive rotational patterns of movement^31^, such as the curvilinear path of travel exhibited by NL3 knockout mice after morphine exposure. D1-MSNs are also a critical site for structural and functional plasticity underlying persistent psychomotor sensitization^32,33,^ suggesting NL3 in these cells could be necessary for the full expression of morphine sensitization.

We began by crossing NL3^fl/fl^ dams with sires from a BAC transgenic line expressing Cre-recombinase under the transcriptional control of the Drd1 dopamine receptor^22^, to achieve selective conditional knockout of NL3 from D1-MSNs (Figure 2A) as previously described^12^. After acute morphine exposure (Figure 2B), all mice exhibited a dose-dependent increase in locomotor activity, as indicated by a main effect of Dose (F_1.509,34.70_=128.3, p<0.0001). However, there was no main effect of Genotype (F_1,23_<1) or interaction (F_3,69_<1), indicating similar distance traveled in both genotypes after acute morphine exposure. Contrary to our expectation, NL3^fl/y^;D1-Cre mice exhibited normal development of psychomotor sensitization after chronic morphine exposure (Figure 2C). Analysis of these data revealed a main effect of Day (F_1,23_=90.47, p<0.0001), but no main effect of Genotype (F_1,23_<1) or interaction (F_1,23_=2.091, p=0.16). The similar degree of sensitization was also apparent in analysis of the change in distance traveled between Day 1 and Day 7 for each individual animal (Figure 2D). Both genotypes also showed a similar response to morphine challenge (Figure 2E), with a main effect of Dose (F_1.448,27.52_=78.14), but no main effect of Genotype (F_1,19_<1) or interaction (F_3,57_<1).

There was no significant difference between genotypes in locomotor topography prior to acute morphine exposure (Figure 2F) (t_24_<1), and a non-significant trend prior to morphine challenge (Figure 2G) (t_20_=2.02, p=0.057). However, there was a Genotype x Dose interaction after morphine challenge (F_1.94,38.88_=3.94, p=0.029), with significant differences between genotypes after challenge with 2 and 6.32 mg/kg morphine. After acute morphine, there was also a non-significant trend toward a main effect of Genotype (F_1,24_=3.86, p=0.061). After morphine administration, NL3^fl/y^;D1-Cre also showed significantly higher absolute angular acceleration (Table S4). NL3^fl/y^;D1-Cre mice thus show a change in locomotor topography after morphine exposure, but no change in psychomotor sensitization, providing further evidence for a dissociation between these phenotypes.

### Conditional NL3 deletion from D2-MSNs does not affect behavioral responses to morphine

Since conditional NL3 deletion from D1-MSNs did not recapitulate the attenuation of psychomotor sensitization seen in NL3 constitutive knockouts, we next considered the role of D2-MSNs, the other major neuron type within the NAc and striatum. Opioid exposure causes functional and structural changes in D2-MSNs^33,34,^ which could be mediated by NL3. As previously described1, we crossed NL3^fl/fl^ dams with sires from a BAC transgenic line expressing Cre-recombinase under the transcriptional control of the Adora2a adenosine receptor (A2a-Cre). This genetic strategy allowed us to target D2-MSNs (Figure 3A), which also exhibit enriched expression of Adora2a^35^, while avoiding other cell types that express Drd2^36^.

After acute morphine exposure (Figure 3B), all mice exhibited a dose-dependent increase in locomotor activity, as indicated by a main effect of Dose (F_2.031,46.71_=231.3, p<0.0001). However, there was no main effect of Genotype (F_1,23_<1) or interaction (F_3,69_<1), indicating that distance traveled was similar in both genotypes after acute morphine exposure. NL3^fl/y^;A2a-Cre mice also exhibited normal development of psychomotor sensitization after chronic morphine exposure (Figure 3C). Analysis of these data revealed a main effect of Day (F_1,16_=163.6, p<0.0001), but no main effect of Genotype (F_1,16_<1) or interaction (F_1,16_=3.743, p=0.071), with a similar change in distance traveled between Day 1 and Day 7 (Figure 3D). Both genotypes also showed a similar response to morphine challenge (Figure 3E), with a main effect of Dose (F_2.167,49.83_=224.4, p<0.0001), but no main effect of Genotype (F_1,23_<1) or interaction (F_3,69_<1). Finally, NL3^fl/y^;A2a-Cre mice exhibited normal locomotor topography before and after both acute morphine exposure (Figure 3F) and morphine challenge (Figure 3G). These results are consistent with a lower level of NL3 expression by D2-MSNs^12^, and suggest this neuronal population is not involved in the altered behavioral responses to morphine exhibited by NL3 constitutive knockout mice.

### Conditional NL3 deletion from dopamine neurons attenuates psychomotor sensitization without affecting locomotor topography

Drug-evoked adaptations in the nucleus accumbens contribute to the long-lasting expression of psychomotor sensitization, but the induction of sensitization involves drug action in the VTA, which contains dopamine neurons that form the mesolimbic projection^19,20.^ Previous studies have shown that NL3 depletion alters the physiology of dopamine neurons, and NL3 expression by these cells is required for normal social behavior^11,37.^ To achieve selective conditional knockout of NL3 from dopamine neurons (Figure 4A), we crossed NL3^fl/fl^ dams with DAT-IRES-Cre (DAT^Cre/wt^) sires^24^. This genetic strategy allowed us to selectively target dopamine neurons while avoiding non-dopaminergic cells^38^.

After acute morphine exposure (Figure 4B), all mice exhibited a dose-dependent increase in locomotor activity, as indicated by a main effect of Dose (F_1.713,29.12_=224.1, p<0.0001). However, there was no main effect of Genotype (F_1,17_<1) or interaction (F_3,51_<1), indicating similar distance travelled in both genotypes after acute morphine exposure. After chronic morphine exposure (Figure 4C), there was a main effect of Day (F_1,17_=152.1, p<0.0001) and a significant Day x Genotype interaction (F_1,17_=5.986, p=0.026). This interaction indicated an attenuation of sensitization in NL3^fl/y^;DAT^Cre/wt^ mice, which was apparent after computing the change in distance traveled between Day 1 and Day 7 for each individual animal (t_13_=2.447, p=0.026) (Figure 4D). This reduced magnitude of sensitization was also evident after morphine challenge (Figure 4E), with a main effect of Dose (F_1.730,68_=220.0, p<0.0001) and significant Dose × Genotype interaction (F_3,51_=3.75, p=0.016). Distance travelled after a challenge injection of 20 mg/kg morphine was significantly reduced in NL3^fl/y^;DAT^Cre/wt^ mutants versus NL3^fl/y^ controls (p=0.032). Conditional genetic deletion of NL3 from dopamine neurons thus recapitulated the diminished psychomotor sensitization seen after chronic morphine exposure in constitutive NL3 knockout mice. Finally, NL3^fl/y^;DAT^Cre/wt^ mutants mice exhibited a normal path of travel before and after both acute morphine exposure (Figure 4F) and morphine challenge (Figure 4G), demonstrating a double-dissociation in the neuronal populations where NL3 regulates psychomotor sensitization and locomotor topography after morphine exposure.

### Photometric correlate of morphine sensitization is absent in VTA dopamine neurons lacking NL3

Opioid administration excites dopamine neurons through a disinhibitory mechanism, involving activation of opioid receptors on inhibitory synaptic inputs^2^. The influence of opioid administration on dopamine neuron activity also changes over the course of chronic opioid exposure^39^. This led us to ask whether NL3 expression shapes the response of dopamine neurons over the course of chronic morphine administration *in vivo*. We used fiber photometry to conduct longitudinal measurements of dopamine neuron activity^40^. To compare these measurements in presence and absence of NL3 expression in dopamine neurons, we bred homozygous DAT^Cre/Cre^ sires with heterozygous NL3^fl/+^ dams, so that male offspring were DAT^Cre/wt^ heterozygotes with or without a floxed NL3 allele (NL3^fl/y^ or NL3^+/y^) (Figure 5A). All mice received unilateral stereotaxic injection of an adeno-associated viral vector encoding Cre-dependent jGCaMP8m (AAV9-hSyn-Flex-jGCaMP8m) into the VTA, followed by implantation of an optical fiber just above the site of virus injection (Figure 5B). Post-mortem histological analysis confirmed robust expression of jGaMP8m in the midbrain (Figure 5C), with anatomical placement of fiber-optic implants in the VTA of both genotypes (Figure 5D).

To record the response of dopamine neurons to morphine administration, we delivered blue light (470 nm) to monitor calcium-dependent changes in jGCaMP8m fluorescence, and purple light (405 nm) as an isosbestic control to correct for bleaching and movement artifact (Figure 5E). At the end of each experiment, to confirm that the fluorescent signal correlated with the activity of dopamine neurons, we administered apomorphine (5 mg/kg) to activate D2-autoreceptors located on the somatodendritic compartment of VTA dopamine neurons^29^. As expected, we observed a consistent decrease in fluorescent signal following apomorphine administration in mice of both genotypes (Figure 5F).

We recorded GCaMP signal on Day 1 and 7 of morphine exposure to measure the response of VTA dopamine neurons (Figure 5G). On the intervening Day 2 and Day 6, we measured distance travelled in the open field after morphine administration (Figure 5H). We again observed a non-significant trend toward attenuation of sensitization in NL3^fl/y^;DAT^Cre/wt^ mice, which was apparent after computing the change in distance traveled for each individual animal (t_12_=2.01, p=0.067) (Figure 5I). Control animals showed a modest increase in ΔF/F following morphine administration on Day 1, which tended to increase after 7 days of morphine (example trace Figure 5J, quantified in Figure 5L; t_8_=2.02, p=0.078), providing a putative photometric correlate of sensitization in VTA dopamine neurons. In contrast, although NL3^fl/y^;DAT^Cre/wt^ animals showed an increase in GCaMP fluorescence on Day 1, this signal did not increase with repeated morphine administration, and in fact was numerically lower on Day 7. This pattern was reflected in a significant Day x Genotype interaction (F_1,20_=5.34, p=0.032) and a significant difference between genotypes in the change in fluorescent response to morphine across repeated exposure (t_20_=2.24, p=0.032) (example trace Figure 5K, quantified in Figure 5M). This result suggests that loss of NL3 expression leads to aberrant changes in the response of dopamine neurons to repeated daily morphine treatment.

### VTA dopamine neurons lacking NL3 exhibit differential synaptic inhibition

Opioid administration excites dopamine neurons by activating opioid receptors on inhibitory synaptic inputs^2^, and NL3 mutations can alter the function of numerous inhibitory synapses in the brain^12,21.^ To assess inhibitory synaptic function in dopamine neurons, we again generated DAT^Cre/wt^ mice with or without a floxed NL3 allele (Figure 5A). All mice received bilateral stereotaxic injection of an adeno-associated viral vector encoding Cre-dependent eYFP (AAV9-EF1a-DIO-eYFP) into the VTA (Figure 6A). This allowed us to identify dopamine neurons in acute brain slices based on yellow fluorescence and perform whole-cell patch-clamp recordings of spontaneous inhibitory postsynaptic currents (sIPSCs) in the presence of glutamate receptor antagonists (Figure 6B). Similar to previous experiments, these mice received seven daily injections of morphine (20 mg/kg) or saline, and acute brain slices were prepared 24 hours after the final injection (Figure 6C).

Analysis of sIPSC amplitude revealed a significant Genotype x Treatment interaction (F_1,16_=6.94, p=0.018) (Figure 6D). In control groups receiving saline injections, sIPSC amplitude was decreased in NL3^fl/y^ mice compared to NL3^+/y^ littermates (p=0.026). This difference was no longer observed after morphine treatment, which significantly increased sIPSC amplitude in NL3^fl/y^ mice relative to the saline control group for this genotype (p=0.018). Synaptic inhibition onto dopamine neurons is thus enhanced by chronic morphine exposure in NL3^fl/y^;DAT^Cre/wt^ mice, which may explain why they exhibit an attenuation of both psychomotor sensitization (Figure 4) and corresponding fiber photometry signals (Figure 5). There were no significant changes in sIPSC frequency in any group (Figure 6E), consistent with prior evidence that morphine injection does not change basal probability of GABA release onto VTA dopamine neurons^41^. Together, these data suggest that loss of NL3 expression fundamentally alters the neurobiological adaptations produced by chronic morphine exposure in VTA dopamine neurons and attenuates the increased dopaminergic sensitivity to morphine normally observed following repeated daily exposure.

## DISCUSSION

Chronic drug exposure causes functional and structural remodeling of synapses in the mesolimbic dopamine system, which could be mediated by synaptic cell adhesion molecules like NL3. Psychomotor sensitization represents a durable enhancement of behavioral sensitivity to morphine that is likely related to synaptic remodeling in the mesolimbic dopamine system. In the absence of NL3, we find that these neuroadaptations appear weakened, leading to attenuation of sensitization. Constitutive NL3 knockout mice exhibit a persistent reduction in the magnitude of psychomotor sensitization caused by intermittent morphine exposure but have a normal psychomotor response to acute morphine. Antinociceptive effects of acute morphine exposure and development of antinociceptive tolerance were also normal in NL3 knockout mice. Due to the X-linkage of the *Nlgn3* gene, all of these analyses were conducted in male mice, which is a key limitation of the present study. We are now in the process of comprehensively analyzing behavioral responses to morphine in female NL3 mutants, using multiple breeding strategies to generate NL3^+/-^ mutants with either NL3^+/+^ or NL3^-/-^ littermates^42^.

We noted that the spatial pattern of locomotor activity, which we refer to as locomotor topography^16^, was altered in NL3 knockout mice after both acute and chronic morphine exposure. Mutant mice exhibited higher angular acceleration and a more curvilinear path of travel after morphine injection, bypassing corners of the chamber and instead traveling through more central areas. While altered travel through the center of an open field arena is often considered anxiety-like behavior in rodents, this may not be generally true for C57Bl/6J mice^43^, especially following morphine exposure^44^. Instead, the locomotor topography phenotype we observe could be related to rotational bias previously reported in NL3 knockout mice^12^, although it was only observed after morphine exposure and not saline injection. A similar pattern has been reported after cocaine (but not saline) injection in mice with a mutation in the serotonin transporter^45^. Rotational phenotypes may be exaggerated when elevated levels of extracellular dopamine activate D1 dopamine receptors^46^. In the NAc of NL3 knockout mice, D1-MSNs have reduced inhibitory synaptic input^12^, and may thus be primed to drive rotational behavior in the presence of elevated dopamine^31^.

Consistent with this hypothesis, conditional knockout of NL3 from D1-MSNs led to a change in locomotor topography after morphine exposure. Previous work suggests this phenotype represents a perseverative pattern of locomotion^46^, and could thus reflect an enhancement of repetitive or stereotyped behavior, which has been linked to NL3 expression in D1-MSNs^12^. However, development of psychomotor sensitization was not affected by conditional genetic knockout of NL3 using either Drd1-Cre (which affects D1-MSNs as well as cells outside the striatum^23^) or Adora2a-Cre. In contrast, conditional knockout of NL3 from dopamine neurons led to an attenuation of psychomotor sensitization after chronic morphine exposure, without affecting locomotor topography. Our results thus reveal a clear double-dissociation in the cell types where NL3 expression is normally required for these two behavioral responses.

Both constitutive NL3 knockout and conditional NL3 deletion from dopamine neurons substantially attenuated psychomotor sensitization, which was evident both during daily morphine injection and following a prolonged drug-free period. This sensitization phenotype was not observed after conditional NL3 deletion from D1-MSNs or D2-MSNs, and thus does not likely involve MSNs in either the NAc or dorsal striatum. Effects on sensitization were robust but incomplete, as mice still showed a diminished increase in distance travelled with repeated daily morphine injection. This could arise from compensatory changes in the absence of NL3 expression, including compensation from other neuroligin family members, since some phenotypes of neuroligin knockout mice are amplified when multiple family members are deleted^47^. Our finding that NL3 is required for full expression of sensitization after chronic morphine treatment implies that this postsynaptic cell-adhesion molecule is an important arbiter of the neuroplasticity induced by repeated morphine. It will be important for future studies to expand upon this analysis by studying chronic treatment with synthetic opioids and other drugs of abuse.

To manipulate NL3 expression by dopamine neurons, we used a DAT-IRES-Cre mouse line^24^. Alterations in locomotor activity have been reported in this mouse line^48^, but not consistently observed^49^. In our initial behavioral studies (Figure 4), any effect of the DAT-IRES-Cre genotype was potentially confounded with NL3 deletion from dopamine neurons. However, any general impact of the DAT-IRES-Cre genotype on locomotion should have been evident during the acute phase of morphine exposure, when the behavior of both genotypes was quite similar. Furthermore, we generated mice for fiber photometry analysis (Figure 5) using a different breeding strategy in which all offspring had the DAT-IRES-Cre genotype. The development of psychomotor sensitization was attenuated in both experiments regardless of breeding strategy, providing further evidence that our results are not confounded by the DAT-IRES-Cre genotype. We used DAT-IRES-Cre mice to selectively target dopamine neurons while avoiding non-dopaminergic cells^38^, but it is important to note that DAT expression is not uniformly high in all dopamine neurons. Lower DAT expression is observed in mesocortical dopamine neurons^50^, so our results may be more closely tied to deletion of NL3 from mesolimbic dopamine neurons.

We directly assessed the function of VTA dopamine neurons using fiber photometry and whole-cell patch-clamp electrophysiology, and found that the impact of chronic morphine administration was either prevented or fundamentally altered in the absence of NL3 expression. After repeated daily morphine injection, the increased calcium signal normally detected in VTA dopamine neurons was not observed after NL3 conditional knockout. This is likely related to an increase in the strength of inhibitory synaptic input onto dopamine neurons that lack NL3 expression following chronic morphine exposure. Surprisingly, that strength of inhibitory synaptic input onto VTA dopamine neurons lacking NL3 was also reduced in saline-treated control mice, relative to the saline control group with normal NL3 expression. However, this reduction of basal synaptic inhibition did not lead to a change in the acute psychomotor response to morphine. This could be due to other cellular or synaptic effects of NL3 deletion, such as increased synaptic incorporation of GluA2-lacking AMPA receptors^11^, which offset the reduction of synaptic inhibition under baseline conditions.

NL3 deletion from VTA dopamine neurons has previously been shown to alter social behavior^11^, and genetic mutations in NL3 have been linked to autism spectrum disorder (ASD) in human patients^7^. While genetic knockout of NL3 produces forms of synaptic dysfunction that resemble the effects of chronic exposure to drugs of abuse^13-15^, we found that NL3 mutant mice were less sensitive to the behavioral impact of chronic morphine exposure. This raises the possibility that drug-evoked forms of synaptic plasticity are occluded by genetic deletion of NL3. Our data also suggest that VTA dopamine neurons exhibit aberrant forms of morphine-evoked synaptic plasticity in the absence of NL3, which may further counteract the effects of addictive drugs. These results are intriguing, given the overlapping circuits implicated in substance use disorders and ASD^51^, and an evolving understanding of the complex ways in which ASD diagnosis affects clinical risk for substance use disorders^52^. They suggest NL3 plays a role in both addiction-related synaptic plasticity and other neuropsychiatric conditions.

## ACKNOWLEDGEMENTS

This research was supported by the University of Minnesota’s MnDRIVE (Minnesota’s Discovery, Research, and Innovation Economy) initiative, as well as NIH grants F30 DA007234 (DDB), T32 DA052109 (PGM, DDB), F30 MH124404 (AK), R00 DA037279 (PER), and R01 DA048946 (PER). The University of Minnesota MnDRIVE Optogenetics Core provided technical support for fiber photometry experiments. Some of the viral vectors used in this study were generated by the University of Minnesota Viral Vector and Cloning Core. We thank Marc Pisansky for technical assistance; Mark Geyer and Richard Sharp for assistance with analysis of the spatial scaling exponent; and Emilia Lefevre, Joshua Melander, Robert Meisel, Bailey Remmers, Carlee Toddes, and Brian Trieu for stimulating discussions. The authors have no conflicts of interest to declare.

## AUTHOR CONTRIBUTIONS

DDB, PER, and PGM were responsible for the study concept and experimental design. DDB and BJS contributed to acquisition of behavioral data. CLR conducted electrophysiology experiments. DDB conducted virus injection and fiber photometry. DDB, CLR, AK, MAM, and PER completed data analysis and figure preparation. DDB and PER drafted the manuscript. CLR and PGM provided critical revision of the manuscript for scientific content. All authors critically reviewed and approved the final version for publication.

