## Supporting information for "Neuroligin-3 in Dopaminergic Circuits Promotes Behavioral and Neurobiological Adaptations to Chronic Morphine Exposure"

### SUPPLEMENTARY METHODS

#### Gene Expression

Quantitative RT-PCR was performed on tissue punches containing the dorsal and ventral striatum, as previously described[1,2]. Tissue was frozen on dry ice and stored at -80°C. RNA was extracted and isolated using the RNeasy Mini Kit (Qiagen) according to manufacturer instructions. A NanoDrop One microvolume spectrophotometer (Thermo Fisher Scientific, Waltham, MA) was used to measure RNA concentration and verify that samples had A260/A280 purity ratio ≥ 2. Reverse transcription was performed using Superscript III (Invitrogen, Eugene, OR). For each sample, duplicate cDNA reactions and subsequent qPCR reactions were conducted in tandem on both samples. Mouse β-actin mRNA was used as the endogenous control, with primer detection sequences of 5'—GAC GGC CAG GTC ATC ACA T—3' and 5'—CCA CCG ATC CAC ACA GAG TA—3'. Primer sequences for detection of NL3 were forward 5'—CAC TGT CTC GGA TTG CTT CA—3' and reverse 5'—TTC CCA GGG CAA TAC AGT CTC—3'. Quantitative RT-PCR using SYBR green (BioRad, Hercules, CA) was carried out with a Lightcycler 480 II (Roche, Basel, Switzerland) using the following cycle parameters: 1 x (30 sec @ 95°C), 35 x (5 sec @ 95°C followed by 30 sec @ 60°C). Results were analyzed by comparing the C(t) values of the treatments tested using the ΔΔC(t) method. Expression values of target genes were first normalized to expression of β-actin. The mean of cDNA replicate reactions was used to quantify the relative target gene expression.

#### Thermal Antinociception

Thermal antinociception was tested in a plexiglass cylinder on a 55°C hot plate (IITC Life Scientific), 15 minutes after morphine administration and just prior to placing animals in open-field chambers for morphine and saline locomotor testing[1]. Percent maximum possible antinociceptive effect (%MPE) was calculated as the percentage difference between the measured nociceptive response latency following morphine administration and baseline nociceptive response latency following saline administration divided by the difference between the maximum response latency and baseline response latency. Nociceptive responses included jumping, hind paw shaking, and hind paw licking. Maximum response latency was defined as 30 sec; animals that failed to respond after 30 sec were removed for the apparatus to avoid tissue damage.

**Measures of Locomotor Activity and Topography**

We tested open-field locomotor activity for 60 minutes in a clear plexiglass arena (ENV-510, Med Associates) within a sound-attenuating chamber. The center zone of the open field arena was defined based on the central 75% in each dimension. To measure angular acceleration, we used raw tracking data to calculate the position of the mouse every 200 ms, which represented a smoothed value of four individual measurements collected at 20 Hz. This smoothing step decreased the granularity and quantal nature of position estimation with infrared photobeams in Activity Monitor software. The moment-to-moment change in x and y position were used to calculate the orientation of motion using the arctangent, which was then unwrapped to prevent circular transitions[3]. These values were used to calculate moment-to-moment changes in the orientation of motion (angular velocity) by differentiation, followed by a second differentiation step to calculate angular acceleration (i.e., change in angular velocity). For analysis, we calculated the average absolute value of angular acceleration for individual mice and individual test sessions, to collapse across acceleration in different directions and focus on the overall magnitude of acceleration. The spatial scaling exponent was calculated as previously described[4].

**Viral Vectors and Stereotaxic Surgery**

The plasmid encoding syn-Flex-jGCaMP8m (Addgene plasmid #162378) was a gift from GENIE Project[5], and packaged in AAV9 by the University of Minnesota Viral Vector and Cloning

Core. The plasmid pAAV-EF1a-DIO-eYFP (Addgene viral prep #27056-AAV9) was a gift from Karl Deisseroth. Intracranial virus injection and optical fiber implantation were performed as previously described[6]. Briefly, mice were anesthetized with a ketamine-xylazine cocktail (100:10 mg/kg), and a burr hole was drilled above target coordinates for the ventral tegmental area (AP -2.9, ML +0.04; DV -4.5). A 33-gauge Hamilton syringe containing the viral solution was lowered to these target coordinates using a stereotaxic apparatus (David Kopf Instruments, Los Angeles, CA). A volume of 500nL of virus was injected at a rate of 100 nL/min. A 6 mm long, 400 μm diameter fiber-optic cannula with 0.48 NA (Doric Lenses: MFC_400/430-0.48_6.0mm_MF2.5_FLT) was inserted just above the site of viral injection (+0.01 mm DV), before being fixed in place with dual-cure resin (Patterson Dental, Inc.) and anchored in place with two skull screws inserted into the parietal bone.

**Immunohistochemistry**

To confirm virus expression and optic cannula placement, all mice were deeply anaesthetized using a phenytoin/pentobarbitol mixture (Beuthanasia, 200 mg/kg, i.p.), then transcardially perfused with ice-cold 0.1M PBS followed by ice-cold 4% paraformaldehyde in 0.1M PBS. Brains were fixed overnight in 4% paraformaldehyde in 0.1M PBS then sliced at 50 μm thickness using a vibratome (Leica VT1000S). Free-floating coronal sections containing the nucleus accumbens were incubated for 3 hours in blocking solution containing 0.2% Triton-X, 2% normal horse serum, and 0.05% Tween20. Sections were then incubated for 48 hours with primary antibodies (Invitrogen mouse anti-eGFP 1:1000, Cat. # A-11120), washed three times, and incubated overnight with secondary antibodies (Abcam goat anti-mouse Alexa Fluor 488 IgG 1:1000, Cat. # ab150117). Slides were washed and mounted using ProLong Gold Antifade mountant with DAPI (Life Technologies). Stained tissue sections were imaged on a laser-scanning confocal microscope (model TCS SPE, Leica Microsystems).

**Table S1. Hot plate antinociception in NL3 constitutive knockout mice after acute and chronic morphine.**

| Genotype | $NL3^{+/y}$ | $NL3^{-/y}$ | *Statistics* |
|---|---|---|---|
| Day 1 | 96.89 ± 2.31 | 85.17 ± 6.42 | $t_{26} = 1.82; p = 0.081$ |
| Day 7 | 50.57 ± 7.24 | 41.05 ± 3.77 | $t_{26} = 1.11; p = 0.28$ |

All data are presented as mean +/- SEM, in units of % maximum possible effect (MPE).

**Table S2. Absolute angular acceleration of locomotor responses to morphine in NL3 constitutive knockout mice.**

| Morphine Dose (mg/kg) | Acute Morphine | | Morphine Challenge | | Genotype Main Effect |
|---|---|---|---|---|---|
| | $NL3^{+/y}$ | $NL3^{-/y}$ | $NL3^{+/y}$ | $NL3^{-/y}$ | |
| 2 | 35.5 ± 1.4 | 39.7 ± 1.0 | 35.6 ± 1.6 | 39.3 ± 1.0 | $F_{1,35} = 9.61$, $p = 0.004$ |
| 6.32 | 29.5 ± 1.4 | 30.1 ± 1.4 | 22.1 ± 2.3 | 27.5 ± 1.0 | |
| 20 | 15.9 ± 1.2 | 17.1 ± 1.4 | 13.1 ± 0.7 | 15.2 ± 0.6 | |

All data are presented as mean +/- SEM, in units of radians per $s^2$.

**Table S3. Spatial scaling exponent of locomotor responses to morphine in NL3 constitutive knockout mice.**

| Morphine Dose (mg/kg) | Acute Morphine | | Morphine Challenge | | Genotype Main Effect |
|---|---|---|---|---|---|
| | $NL3^{+/y}$ | $NL3^{-/y}$ | $NL3^{+/y}$ | $NL3^{-/y}$ | |
| 2 | 1.36 ± 0.04 | 1.36 ± 0.03 | 1.30 ± 0.03 | 1.33 ± 0.02 | $F_{1,35} < 1$ |
| 6.32 | 1.30 ± 0.03 | 1.28 ± 0.03 | 1.27 ± 0.03 | 1.34 ± 0.06 | |
| 20 | 1.20 ± 0.02 | 1.21 ± 0.02 | 1.22 ± 0.03 | 1.22 ± 0.02 | |

All data are presented as mean +/- SEM of the spatial scaling exponent (*d*).

**Table S4. Absolute angular acceleration of locomotor responses to morphine after conditional knockout of NL3 via genetic cross to Drd1-Cre.**

| Morphine Dose (mg/kg) | Acute Morphine | | Morphine Challenge | | Genotype Main Effect |
|---|---|---|---|---|---|
| | $NL3^{fl/y}$ | +D1-Cre | $NL3^{fl/y}$ | +D1-Cre | |
| 2 | 36.0 ± 0.9 | 36.0 ± 1.6 | 34.3 ± 0.9 | 35.1 ± 1.1 | $F_{1,43} = 5.02$, $p = 0.030$ |
| 6.32 | 18.0 ± 1.7 | 22.8 ± 1.7 | 18.3 ± 1.0 | 20.7 ± 1.5 | |
| 20 | 11.0 ± 0.9 | 12.1 ± 1.3 | 10.5 ± 0.7 | 12.5 ± 1.2 | |

All data are presented as mean +/- SEM, in units of radians per $s^2$.
